# VrySure: A Multi-Task AI Scientific Fraud Detection Platform for Identifying Manipulated and AI-Generated Biomedical Research Images

**DOI:** 10.64898/2026.06.10.731492

**Authors:** Jiacheng Sun, Bingrui Li, Raghu Kalluri

## Abstract

Integrity of scientific data is critical in biomedical research, where images often serve as primary evidence for experimental observations and conclusions. Advances in image-editing technologies and generative artificial intelligence (AI) have increased the accessibility and realism of visual manipulation, making detection through manual review increasingly challenging. To empower our laboratory researchers to continuously monitor and uphold scientific rigor and data integrity, and serve the global scientific community, we developed VrySure, an easy-to-deploy, AI-driven multi-task platform for automated image-integrity screening in biomedical research. VrySure integrates four detection modules: cross-image transformation detection, within-image copy-move detection, splicing detection in blot and gel images, and AI-generated image detection. The system identifies potentially manipulated images and, when possible, localizes suspicious regions using bounding-box outputs to support downstream verification. To support development and evaluation, we constructed task-specific datasets by combining public biomedical image resources, curated manipulated examples, and synthetic images generated by multiple generative AI systems. We evaluated VrySure using region-level F1 score, recall, precision, false negative rate (FNR), and false discovery rate (FDR) across multiple manipulation categories and compared its performance with two commonly used commercial image-integrity screening platforms under a predefined benchmark protocol. Under the tested conditions, VrySure achieved a higher F1 score and recall, lower FNR, and maintained a low FDR for within-image copy-move detection, splicing detection, and AI-generated image detection, while showing comparable performance in transformation detection. Beyond automated screening, VrySure is designed to support source-data comparison and evidence-based assessment in scientific integrity investigations. By integrating multiple detection capabilities into a unified and scalable workflow, VrySure provides a practical framework to improve the efficiency and consistency of image-integrity screening in biomedical research.

## Introduction

Scientific research integrity assessment has long been a critical challenge in biomedical research, and recent advances in image-editing and generative AI technologies have significantly intensified this problem^1–3^. As manipulation tools become more accessible, realistic and scalable, distinguishing authentic scientific images from altered or synthetically generated ones is increasingly difficult. This growing uncertainty threatens the reliability of published findings and undermines trust in the scientific record. Large-scale screening studies have documented the prevalence of problematic image duplication in biomedical publications, and recent studies demonstrate that generative AI can produce highly realistic scientific images that are difficult to detect by visual inspection or existing forensic tools^3,4^. The scale of modern scientific publishing, combined with the rapid adoption of AI tools in research workflows, further amplifies the urgency of this issue.

Biomedical research relies heavily on image-based evidence to document experimental observations and support scientific conclusions. In many cases, images are not merely illustrative but serve as primary data for interpretation, validation, and reproducibility^1,2^. Although quantitative measurements are essential, they often fail to capture important visual characteristics such as spatial organization, cellular morphology, and staining patterns that are critical to biological interpretation^1,2^. As a result, manipulated or AI-generated images can introduce substantial distortions in scientific understanding.

The consequences of compromised image integrity extend beyond individual publications. At the laboratory level, unreliable image-based conclusions can lead to wasted resources and misguided experimental efforts. At the field level, falsified visual evidence can skew research directions and erode confidence in legitimate findings^4,5^. Broader impacts also arise when flawed publications influence clinical decision-making or are incorporated into downstream knowledge systems, including AI-based tools that synthesize scientific literature. These risks highlight the need for systematic and scalable approaches to image-integrity assessment.

Despite its importance, robust image-integrity screening remains challenging in real-world workflows. Manual inspection is still widely used but is work-intensive, subjective and difficult to scale across large volumes of submissions^4,6,7^. Subtle manipulations may be overlooked, particularly under time constraints, and judgments can vary across reviewers. Post-publication platforms provide important community oversight, but post-publication review alone may not be sufficient to prevent problematic images from entering the scientific record. In this regard, there is an urgent need for systematic pre-publication screening employing user-friendly and uncomplicated AI tools^6^. Existing automated approaches vary in accuracy, usability, and workflow integration across tasks and settings. Therefore, these limitations highlight the need for a more scalable, accurate, and comprehensive solution.

To address these challenges, we invented VrySure, an AI-driven multi-task framework for automated biomedical image-integrity screening. VrySure supports detection across multiple categories of manipulation, including cross-image transformation, within-image copy-move duplication, splicing in blot and gel images, and AI-generated content. By combining complementary detection modules within a unified workflow, VrySure aims to provide scalable and consistent identification of suspicious image regions to support downstream verification. In the following sections, we describe the system design, dataset construction, and benchmarking results, and evaluate VrySure against existing image-integrity detection tools.

### Problem Formulation and Task Definition

We categorize biomedical image-integrity violations into four common manipulation types: transformation, copy-move, splicing, and AI-generated imagery. Transformation refers to reused image content that has been altered through geometric operations such as rotation, flipping, scaling, translation, cropping, or a combination of these operations. Copy-move manipulation involves one or multiple paired duplicating regions within the same image. Splicing refers to combining content from multiple images into a single composite. AI-generated imagery refers to images produced fully or partially by generative models.

Given a set of input images *X* = {*I*_1_, *I*_2_, … , *I*_*m*_}, VrySure aims to determine whether each image contains evidence of manipulation and, when possible, localize the suspicious regions. We define VrySure as a set of detection modules *V* = {*T*, *C*, *S*, *A*}, where *T* denotes transformation detection, *C* denotes copy-move detection, *S* denotes splicing detection, and *A* denotes generative AI image detection. For each input image *I*_*i*_, VrySure returns either no detection or a set of bounding boxes *B*_*i*_ = {*b*_*i*1_, *b*_*i*2_, … , *b*_*in*_}, where each bounding box *b*_*ij*_ = [*x*_*min*_, *y*_*min*_, *x*_*max*_, *y*_*max*_] denotes a region associated with a potential integrity concern. At the manuscript level, the overall output is given by *V*(*X*) = {*B*_1_, *B*_2_, … , *B*_*m*_}, enabling both image-level screening and region-level localization.

Metrics used to evaluate performance were F1 scores, precision, recall, false negative rate (FNR), and false discovery rate (FDR). These metrics were defined as follows: 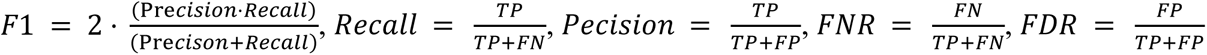 where *TP* denotes the number of true positives, *FP* denotes the number of false positives, and *FN* denotes the number of false negatives.

### System Overview

VrySure is an AI-driven multi-task system designed for automated image-integrity screening in biomedical research. The system integrates a complete workflow from file input to expert review and decision support (**Figure 1**). VrySure accepts common image file types, including JPG, PNG, and TIFF, as well as complete manuscripts in PDF format. After input, the system performs file preprocessing and subimage extraction by parsing the submitted files, extracting individual figures and subimages, and converting them into appropriate formats for downstream analysis.

**Figure 1.**
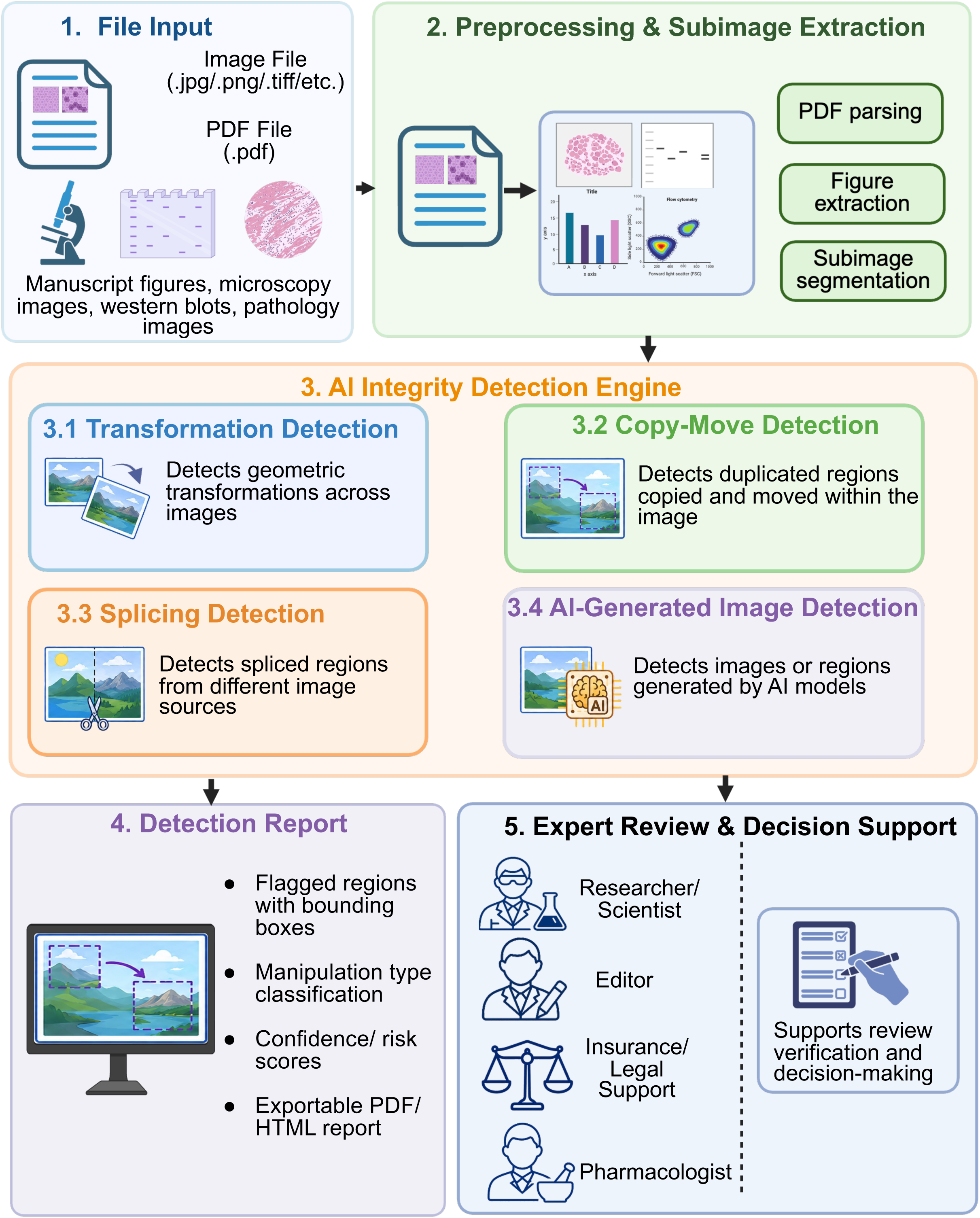
VrySure system overview. The VrySure integrity-checking system consists of five major steps: file input, file preprocessing and subimage extraction, core detection, report generation, and expert review/decision support. VrySure accepts common image file types, including JPG, PNG, and TIFF, as well as complete manuscripts in PDF format. During preprocessing, the system parses the input files, extracts individual images and subimages, and converts them into appropriate formats for downstream analysis. The preprocessed images are then analyzed by four core detection modules: transformation detection, copy-move detection, splicing detection, and AI-generated image detection. These modules identify suspicious regions and generate detection decisions, which are subsequently used to produce a report that marks suspicious areas with bounding boxes, classifies potential manipulation types, and assigns risk scores. The final report supports researchers, journal editors, pharmacologists, insurance professionals, legal professionals, and other stakeholders who require evidence-based image integrity assessment.

The core of VrySure consists of four complementary detection modules addressing distinct classes of image manipulation: cross-image transformation, within-image copy-move duplication, splicing in blot and gel images, and AI-generated content. These modules capture common mechanisms by which visual evidence may be altered or fabricated while preserving a plausible experimental appearance. Transformation, copy-move, and splicing detection focus on identifying reused, duplicated, or composited visual elements, whereas AI-generated image detection targets images synthesized or modified using generative models.

Preprocessed images are forwarded through each detection module to identify suspicious regions and generate detection decisions. These decisions are then used for report generation, in which suspicious areas are marked with bounding boxes, potential manipulation types are classified, and risk scores are assigned. In addition to automated detection, VrySure supports comparison against source data when available, enabling evidence-based verification in disputed cases. This design allows the system not only to flag suspicious regions but also to support downstream expert review, editorial assessment, pharmacological evaluation, insurance review, legal investigation, and other decision-making processes requiring reliable image-integrity evidence.

### Benchmark Evaluation

We evaluated VrySure against two widely used commercial image-integrity tools, Platform A and Platform B, using curated benchmark datasets covering all four manipulation types. Because these tools are not open-source and do not provide full pixel-level outputs, we adopted a region-level evaluation protocol. A prediction was considered a true positive if any detected bounding box overlapped with a ground-truth manipulation region, and a false positive if it only overlapped with a background or non-manipulated region. Platform A and Platform B were evaluated using their respective online interfaces with default or recommended settings, accessed between March and May 2026. No tool-specific parameter tuning was performed. The same input images, formats, and resolutions were used across all platforms. Performance was evaluated using the F1 score, recall, precision, FNR and FDR, and the F1 score was used to summarize the balance of performance between precision and recall.

### Transformation Detection

We evaluated transformation detection using 48 original images and their corresponding transformed versions, grouped into 48 matched pairs along with 1,056 unmatched pairs. Examples of transformation detection are shown in **Figure 2A**. VrySure showed performance comparable to Platform B, with both systems reaching an F1 score of 0.95, and achieved a higher F1 score than Platform A under the evaluated conditions (F1 = 0.70) (**Figure 2B**, **Table 1**). These results suggest that transformation-based duplication can be effectively detected across different tools in this benchmark setting.

**Figure 2.**
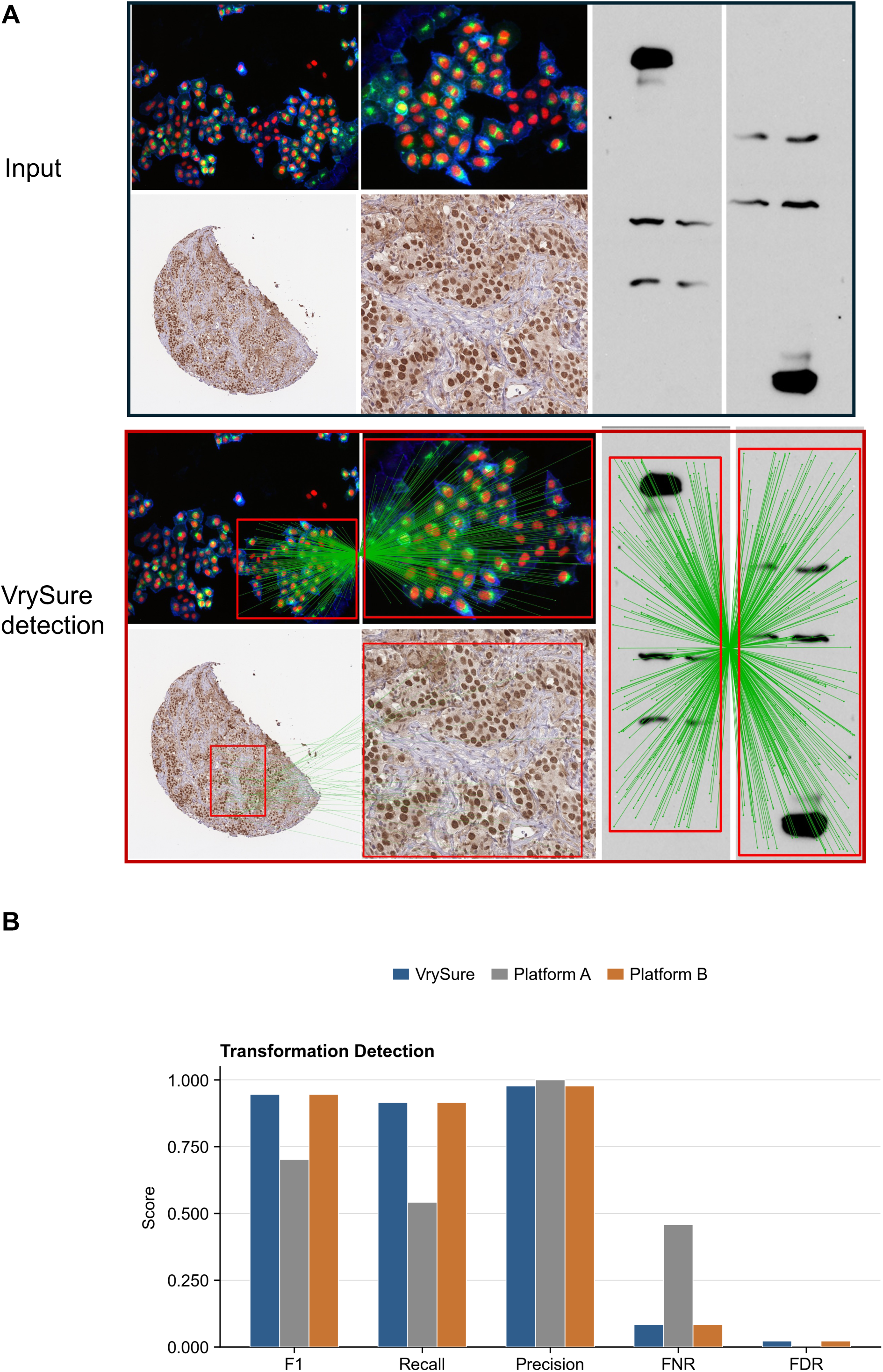
Demonstration and performance of transformation detection. (A) Example of transformation detection. Three input examples are shown on the top panel. In each example, the image on the left is the original image, and the image on the right is the corresponding transformed image. Three output detection examples with connected inlier matches are shown on the bottom panel. The area enclosed by the bounding box indicates the suspicious region. (B) The bar plot shows the F1 scores, recall, precision, false negative rate and false discovery rate for transformation detection across Platform A, Platform B, and VrySure. Results are specific to the evaluated dataset, software versions, input conditions, and benchmark protocol.

**Table 1.** Performance table of transformation detection.

| Platforms | F1 | Recall | Precision | FNR | FDR |
| --- | --- | --- | --- | --- | --- |
| Platform A | 0.703 | 0.542 | **1.000** | 0.458 | **0.000** |
| Platform B | **0.946** | **0.917** | 0.978 | **0.083** | 0.022 |
| VrySure | **0.946** | **0.917** | 0.978 | **0.083** | 0.022 |

### Copy-Move Detection

The copy-move benchmark consisted of 100 images, including 50 manipulated images containing 159 duplicated regions and 50 non-manipulated images. Examples are shown in **Figure 3A**. Under the evaluated conditions, VrySure achieved an F1 score of 0.605, compared with 0.425 for Platform A and 0.262 for Platform B (**Figure 3B**, **Table 2**). Despite these differences, overall performance across all methods remains moderate, reflecting the inherent difficulty of distinguishing duplicated regions from visually similar biological structures.

**Figure 3.**
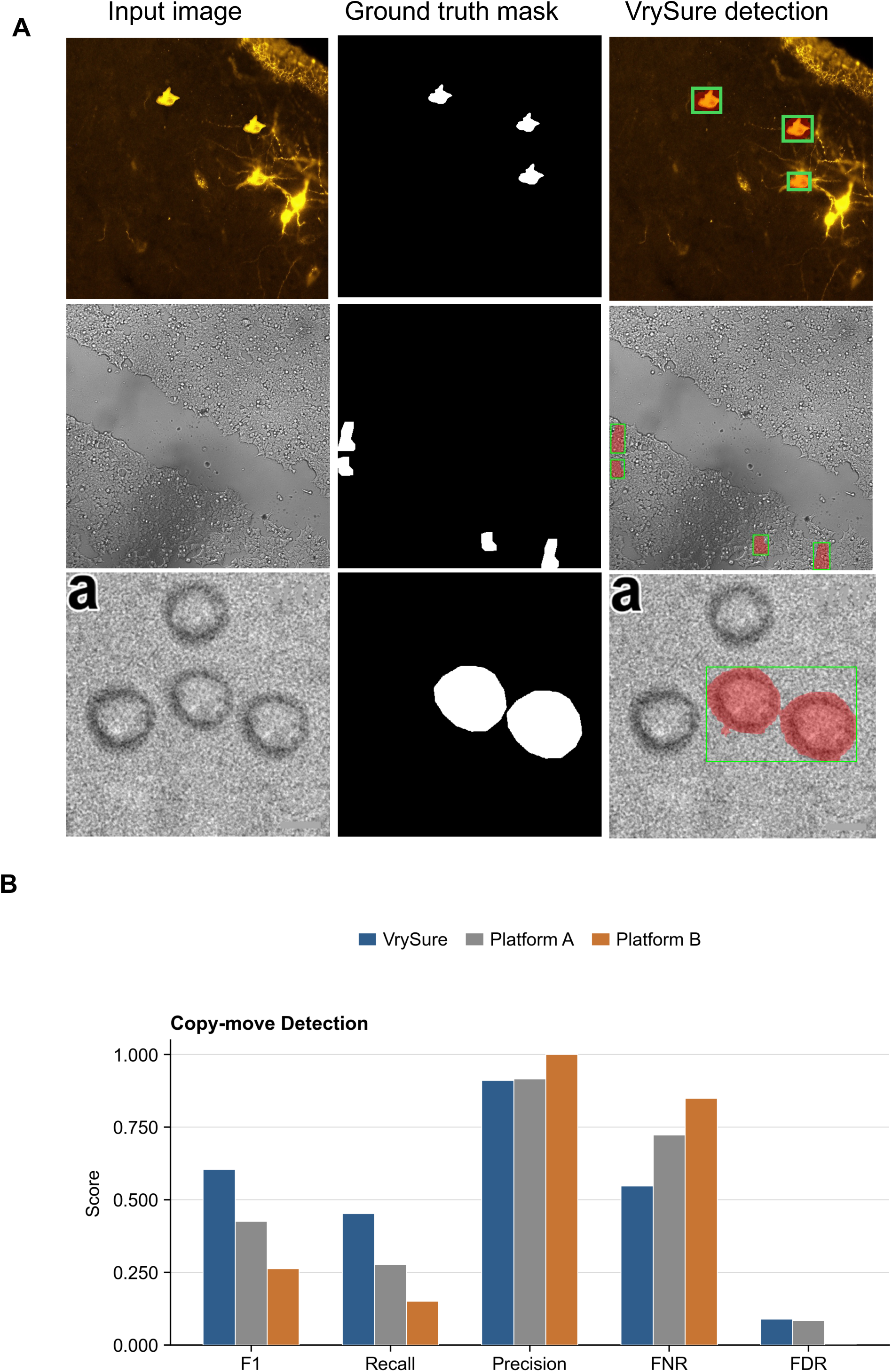
Demonstration and performance of copy-move detection. (A) Example of copy-move detection. The images in the left column are input images containing copy-move regions. The images in the middle column show the ground-truth masks, and the images in the right column show the corresponding detection results. The areas enclosed by the bounding boxes indicate the suspicious regions. Each row represents one image group. (B) The bar plot shows the F1 scores, recall, precision, false negative rate and false discovery rate for copy-move detection across Platform A, Platform B, and VrySure. Results are specific to the evaluated dataset, software versions, input conditions, and benchmark protocol.

**Table 2.** Performance table of copy-move detection.

| Platforms | F1 | Recall | Precision | FNR | FDR |
| --- | --- | --- | --- | --- | --- |
| Platform A | 0.425 | 0.277 | 0.917 | 0.723 | 0.083 |
| Platform B | 0.262 | 0.151 | **1.000** | 0.849 | **0.000** |
| VrySure | **0.605** | **0.453** | 0.911 | **0.547** | 0.089 |

### Splicing Detection

The splicing dataset included 100 images, with 50 manipulated images containing 142 splice boundaries and 50 non-manipulated images. Representative examples are shown in **Figure 4A**. Under the evaluated conditions, VrySure achieved an F1 score of 0.877, compared with 0.185 for Platform A and 0.081 for Platform B (**Figure 4B**, **Table 3**). These differences suggest that task-specific training and localization strategies may contribute to improved detection of splice-related artifacts in this benchmark setting.

**Figure 4.**
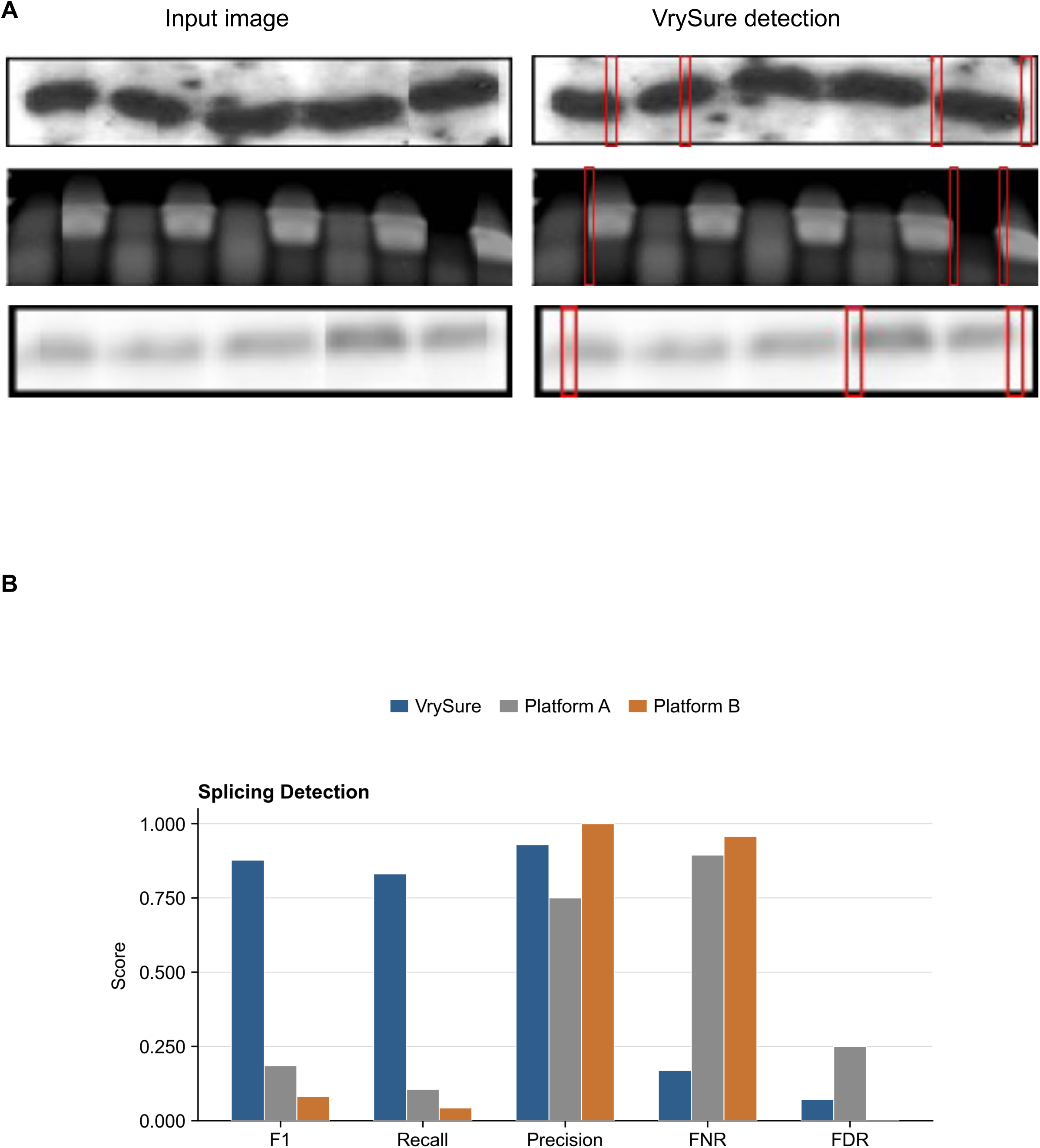
Demonstration and performance of splicing detection. (A) Example of splicing detection. The images on the left are input images, and the images on the right show the detection results. The areas enclosed by the bounding boxes indicate the suspicious regions. (B) The bar plot shows the F1 scores, recall, precision, false negative rate and false discovery rate for splicing detection across Platform A, Platform B, and VrySure. Results are specific to the evaluated dataset, software versions, input conditions, and benchmark protocol.

**Table 3.** Performance table of splicing detection.

| Platforms | F1 | Recall | Precision | FNR | FDR |
| --- | --- | --- | --- | --- | --- |
| Platform A | 0.185 | 0.106 | 0.75 | 0.894 | 0.25 |
| Platform B | 0.081 | 0.042 | **1.000** | 0.958 | **0.000** |
| VrySure | **0.877** | **0.831** | 0.929 | **0.169** | 0.071 |

### AI-Generated Image Detection

The AI-generated image benchmark included 100 images, evenly split between synthetic and authentic images. Synthetic images were generated using multiple generative AI systems (**Figure 5A**). Under the evaluated conditions, VrySure achieved an F1 score of 0.98, compared with 0.33 for Platform A and 0.14 for Platform B (**Figure 5B**, **Table 4**). These results suggest that incorporating dedicated detection strategies may be beneficial for identifying AI-generated content in this benchmark setting as generative models continue to produce increasingly realistic biomedical images.

**Figure 5.**
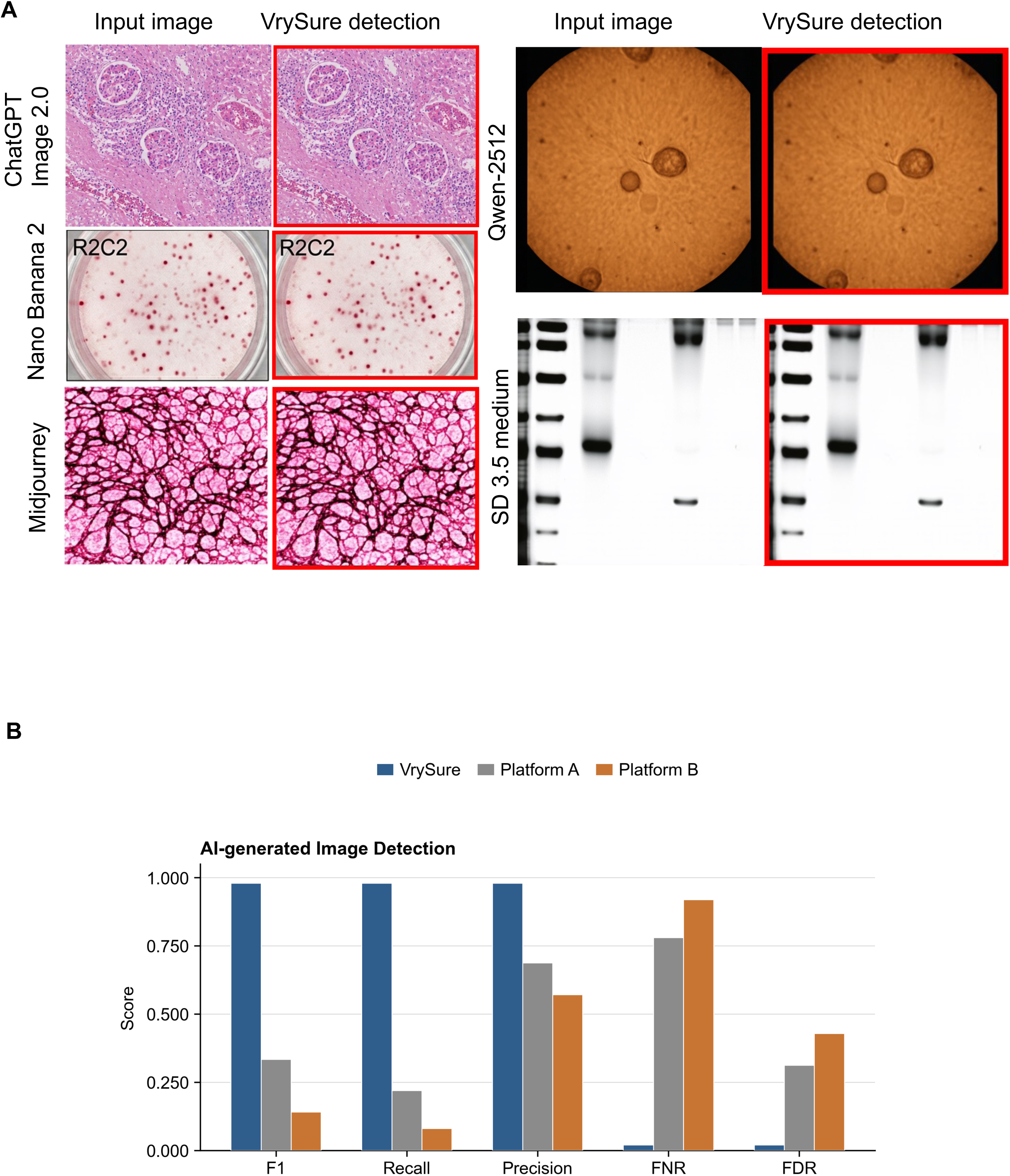
Demonstration and performance of AI-generated image detection. (A) Example of AI-generated image detection. Each input image is paired with its corresponding detection result. The areas enclosed by the bounding box indicate the suspicious regions. The image–result pairs are arranged in two columns: the left column contains three pairs and the right column contains two pairs. From top to bottom in the left column, the pairs correspond to images generated by ChatGPT Image 2.0, Nano Banana 2, and Midjourney. From top to bottom in the right column, the pairs correspond to images generated by Qwen-2512 and Stable Diffusion 3.5 medium. (B) The bar plot shows the F1 scores, recall, precision, false negative rate and false discovery rate for AI-generated image detection across Platform A, Platform B, and VrySure. Results are specific to the evaluated dataset, software versions, input conditions, and benchmark protocol.

**Table 4.** Performance table of AI-generated image detection.

| Platforms | F1 | Recall | Precision | FNR | FDR |
| --- | --- | --- | --- | --- | --- |
| Platform A | 0.333 | 0.220 | 0.688 | 0.780 | 0.313 |
| Platform B | 0.140 | 0.080 | 0.571 | 0.920 | 0.429 |
| VrySure | **0.980** | **0.980** | **0.980** | **0.020** | **0.020** |

## Discussion

Our laboratory provides an AI-driven framework for scalable image-integrity screening in biomedical research, named here as VrySure. By conducting a thorough evaluation of all scientific data in real time as experimental data is documented, during figure generation prior to peer review, the system has the potential to avoid human errors, inadvertent image duplications or purposeful deceit and improve the reliability of research results. VrySure may help reduce the risk that subsequent research is built on potentially unreliable scientific evidence. In addition to automated detection, VrySure is designed to support evidence-based assessment by enabling comparison with source data when available, which may be particularly valuable in resolving disputed scientific integrity cases.

As a scalable system, VrySure also has the potential to be integrated into broader editorial and research-integrity workflows through SaaS deployment or API access. Such integration could accelerate automated screening in agent-based large language model (LLM) review systems and may help improve detection coverage in large publication workflows. However, several limitations remain. First, generative AI models evolve rapidly, and detection models may lag behind newly released image-generation systems. This may create a coverage gap in which images produced by newer models may be more difficult to detect^3^. Second, the rapid iteration of competing generative AI systems may challenge the stability of AI-image detection and increase the cost of model maintenance^3,6^. Third, splicing detection currently relies on identifying evidence such as splice seams, which may be weakened or removed by additional image processing. Unlike duplicated regions, a removed splice seam may leave insufficient visual evidence to determine whether an image has been manipulated. For this reason, raw source data remains essential as the most reliable ground-truth evidence in disputed scientific integrity investigations^2,3^.

In addition, the benchmark results presented in this study are specific to the evaluated datasets, manipulation categories, software versions, and testing protocol and may not fully generalize to all real-world scenarios or alternative workflows. Performance of comparator platforms may vary depending on configuration, input conditions, and future software updates.

Future development will focus on model updating, production deployment, and extensibility. To provide a practical user experience, VrySure should maintain high throughput while balancing computational cost and system performance. The current design separates the detection models into modular services to improve flexibility, but an integrated large-model architecture may also be explored in future work as a trade-off between performance, maintainability, and deployment complexity. These alternatives should be evaluated through systematic testing and user-centered workflow studies.

## Methods

We constructed and used task-specific datasets for training and evaluation instead of relying exclusively on BioFors or other public datasets. This approach was used to support consistent annotation and evaluation. BioFors was created from biomedical research documents and includes real-world suspicious manipulations, but some cases may require additional interpretation of provenance and context^7^. In addition, meaningful benchmarking requires clear annotation of the manipulated regions; without accurate ground-truth areas, performance cannot be interpreted reliably^7,8^. We therefore combined public datasets with curated manipulated and generated examples to reduce dependence on any single data source and to evaluate performance across multiple forms of image manipulation. All benchmark datasets were defined prior to comparative evaluation, and test sets were held out from model development and tuning to prevent data leakage.

The transformation dataset was used primarily for classifier training to determine whether one image is a transformed version of another. Classification was based on image-matching and geometric-consistency metrics, including “n_matches”, “n_inliers”, “inlier_ratio”, “reproj_inliers_rmse”, “reproj_all_rmse”, “reproj_all_median”, “reproj_all_p90”, “rmse_ratio”, “p90_minus_median”, “hull_area_ratio0”, “hull_area_ratio1”, “bbox_area_ratio0”, “bbox_area_ratio1”, “grid_occupancy0”, “grid_occupancy1”, “std_x0”, “std_y0”, “std_x1”, “inlier_centroid_dist_to_center”, “H_cond”, “detA”, “detA_sign”, “projective_mag”, “translation_norm”, “dist_ratio_std”, “dist_ratio_mad”, “angle_diff_std”, “angle_diff_mad”, and “best_flip_id”. The dataset includes both original images and transformed versions with comparisons performed across all possible image pairs. Each transformed image was generated by randomly applying flipping, rotation, translation, scaling, or a combination of these operations, with each transformation type sampled with equal probability. Benchmark evaluation was performed on a held-out subset not used during model training.

The copy-move dataset includes images from D2PRL and Recod.ai/LUC - Scientific Image Forgery Detection/training. To make D2PRL compatible with our model format, we converted its masks into binary masks. In Recod.ai/LUC - Scientific Image Forgery Detection/training, copied regions are provided as separate masks for individual pairs, even when multiple pairs appear in the same image. We therefore merged the separate masks into a single mask for each image and converted the result into the required image-mask format. All mask transformations and merging procedures were applied consistently across the dataset.

The splicing dataset was constructed from blot-gel images in BioFors. We extracted blot-gel images without confirmed fraud and manually removed images with uncertain integrity status. After constructing a clean set of non-manipulated blot-gel images, we generated spliced examples by extracting a random number of 5-pixel-wide strips from the original images and generated additional spliced examples using controlled procedures designed to represent common splice patterns observed in biomedical images. These synthetic manipulations may not fully represent all real-world splicing scenarios.

The generative AI dataset was built in-house. We generated synthetic biomedical images using ChatGPT Image 2.0, Gemini, Stable Diffusion 3.5 medium, Midjourney, and Qwen-image-2512. Images from Stable Diffusion and Qwen-image-2512 were generated using scripts based on Hugging Face instructions. Synthetic biomedical images were generated across several categories, including microscopy, histology, flow cytometry, and gel/blot images. Images from ChatGPT, Gemini, and Midjourney were generated by entering prompts through their client interfaces. To automate part of this process, we developed JavaScript scripts for browser-based generation workflows. To reduce cost and improve efficiency, we generated composite images containing approximately 12 subimages each, then used our subimage extraction algorithm to separate these subimages and treat them as individual dataset samples. Generated images may not fully capture the diversity of outputs from all generative models or real-world usage scenarios.

The same input images, formats, and preprocessing procedures were used across all evaluated tools to ensure consistent comparison.

## Datasets

To train and fine-tune the transformation, copy-move, splicing, and generative AI detection modules, we constructed task-specific datasets for model development and evaluation.

The collected data sources include BioFors^7^, Recod.ai/LUC - Scientific Image Forgery Detection^9^, the Scientific Image Classification Dataset^10^, D2PRL^11^, IDR^12^, and publicly available datasets, as well as internally collected images. The 500 FACS images are from the Scientific Image Classification Dataset^10^.

For AI-generated image detection, we generated synthetic biomedical images using the ChatGPT client, Gemini client, Stable Diffusion 3.5 Medium from Stability AI, Midjourney client, and Qwen-Image-2512. ChatGPT, Gemini, and Stable Diffusion were used to generate all biomedical image categories, including flow cytometry/FACS plots, western blot and gel electrophoresis images, histology and histopathology images, microscopy images, fluorescence microscopy images and cell culture images. Midjourney and Qwen-image-2512 were used to generate biomedical image categories, excluding blot-gel images.

We also regrouped existing data and generated manipulated examples to match the requirements of each detection module. The transformation dataset contains 500 FACS images from the Scientific Image Classification Dataset and all authentic images from Recod.ai/LUC - Scientific Image Forgery Detection/train_images/authentic, together with their transformed versions. The copy-move dataset contains images from D2PRL and Recod.ai/LUC - Scientific Image Forgery Detection/train with corresponding masks. The splicing dataset contains manually filtered non-fraud blot-gel images from BioFors and their manipulated versions. For each dataset, images were split into training (80%) and held-out test (20%) sets, with no overlap between training and evaluation data. This split was applied independently to the transformation dataset, the copy-move dataset with masks, the splicing dataset, and the AI-generated image dataset. Synthetic and curated examples were included to represent specific manipulation types and were combined with real-world images to reduce dependence on any single data source; however, they may not fully capture all real-world variability.

## Acknowledgments

This work was supported by the Sid Richardson Foundation and Kalluri laboratory research funds provided by UT MD Anderson Cancer Center. If anyone is interested in obtaining the operational code for this technology from UT MD Anderson Cancer Center, please contact the corresponding author for relevant information.

